# Targeted photostimulation uncovers circuit motifs supporting short-term memory

**DOI:** 10.1101/623785

**Authors:** Kayvon Daie, Karel Svoboda, Shaul Druckmann

## Abstract

Short-term memory is associated with persistent neural activity without sustained input, arising from the interactions between neurons with short time constants^1,2^. A variety of neural circuit motifs could account for measured neural activity^3–7^. A mechanistic understanding of the neural circuits supporting short-term memory requires probing network connectivity between functionally characterized neurons^8^. We performed targeted photostimulation of small (< 10) groups of neurons, while imaging the response of hundreds of other neurons^9,10^, in anterior-lateral motor cortex (ALM) of mice performing a delayed response task^11^. Mice were instructed with brief auditory stimuli to make directional movements (lick left or lick right), but only after a three second delay epoch. ALM contains neurons with delay epoch activity that is selective for left or right choices. Targeted photostimulation of groups of neurons during the delay epoch allowed us to observe the functional organization of population activity and recurrent interactions underlying short-term memory. These experiments revealed strong coupling between neurons sharing similar selectivity. Brief photostimulation of functionally related neurons produced changes in activity in sparse subpopulations of nearby neurons that persisted for several seconds following stimulus offset, far outlasting the duration of the perturbation. Photostimulation produced behavioral biases that were predictable based on the selectivity of the perturbed neuronal population. These results suggest that ALM contains multiple intercalated modules, consisting of recurrently coupled neurons, that can independently maintain persistent activity.

## Main text

Mice expressing the calcium indicator GCaMP6s^12^ and the light-activated cation channel soma-targeted (ST) ChrimsonR^13,14^ (Extended Data Fig. 1) were trained to discriminate two tones. After a delay epoch lasting three seconds, mice reported the identity of the tone with directional licking (Fig. 1a)^15^. We imaged activity in layer 2/3 (125-250 um deep; typical field of view (FOV), 600 x 600 μm^2^) of the left hemisphere of ALM (8 mice, 84 sessions, 324 trials per session; range, 208 – 441 trials; Extended Data Fig. 2). On average, 49 neurons (range, 14 — 98, 75% CI; 25 % of the imaged population) per FOV were selective during the delay and early response epochs (mean, 26 right- and 23 left-selective, Extended Data Fig. 3, one-tailed t-test, p < 0.05)^16,17^.

**Figure 1:**
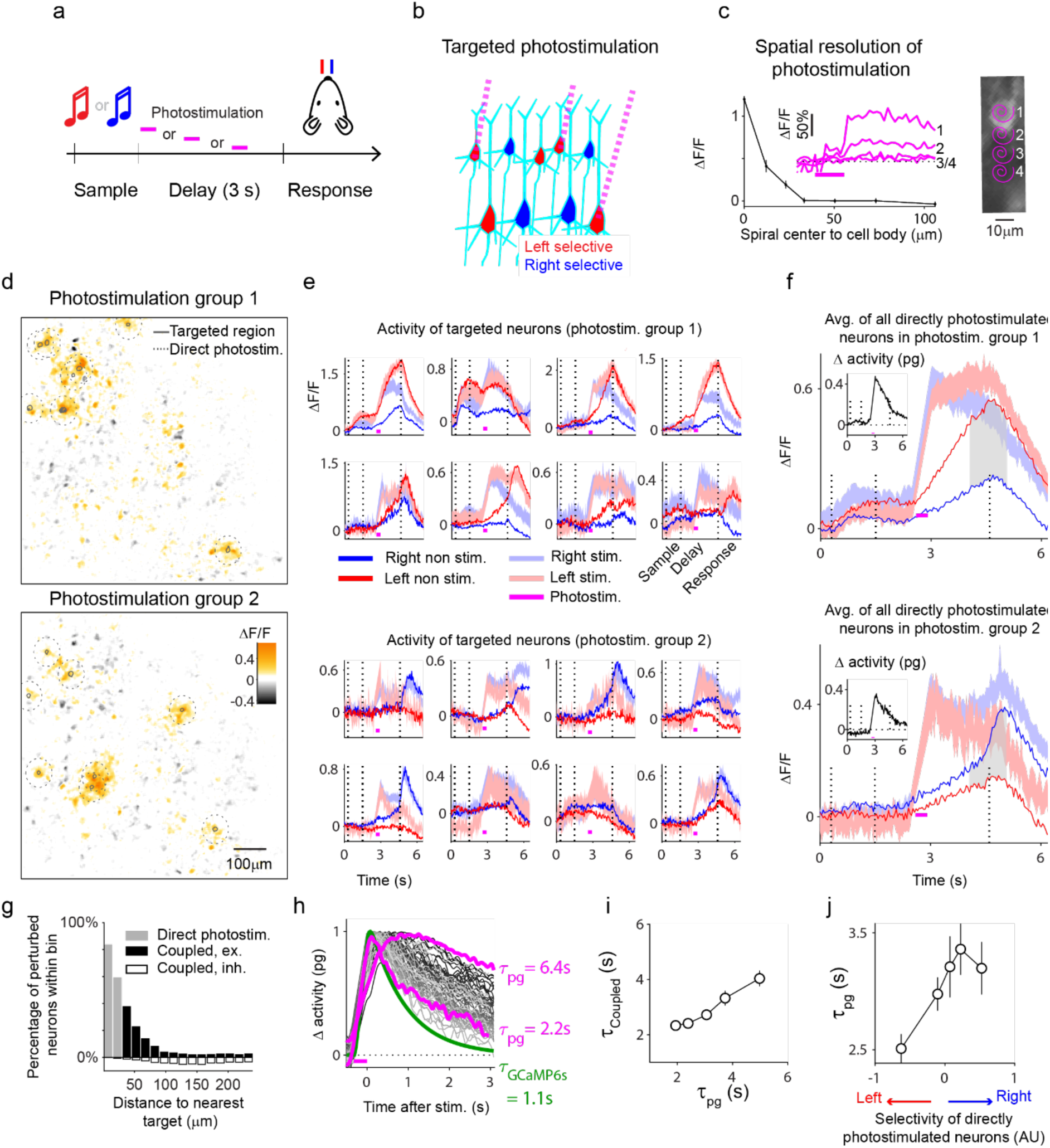
Targeted photostimulation during performance of the delayed-response task. **a**. Task structure. During the sample epoch, mice were instructed by an auditory cue to lick left or lick right for a water reward. Responses were allowed after a 3 seconds long delay epoch. Photostimuli (magenta bars) were delivered on a random subset of trials during the delay epoch. **b**. Targeted photostimulation. Two-photon imaging was used to map selectivity of individual neurons during behavior (lick left, red; lick right, blue). Two-photon photostimulation (magenta dashed lines) was used to activate groups of neurons based on their selectivity. **c.** Measurement of lateral spatial resolution of photostimulation. Left, change in GCaMP6s fluorescence vs. distance between photostimulus (spiral) and imaged neuron (9 neurons). Inset, responses to photostimuli delivered at different distances (μm; 0, 12.5, 25 and 37.5) (magenta bar, photostimulation, 320 ms). Right, example experiment. **d- f.** Top, photostimulation group 1; bottom, photostimulation group 2. **d**, Example photostimulation experiment. Black ROIs indicate targeted neurons. Dashed circles (radius 20 μm) indicate region with neurons that could be receiving significant direct photostimulation. Pixel color indicates the fractional difference in fluorescence between photostimulation and non-photostimulation trials, averaged over 500 ms after the photostimulus. (Excited coupled neurons, 22 in group 1, 23 in group 2; Inhibited coupled neurons, 6 in group 1, 7 in group 2; p < 0.05, one-tailed T-test) **e.** Fluorescence changes (averaged across correct trials) for each of the eight targeted neurons per photostimulation group, on photostimulation (light traces; error shade = s.e.m.) and non-photostimulation (dark traces) trials (trial types: red, left; blue, right). **f.** Responses averaged across all directly photostimulated neurons per photostimulation group. Gray shading, average selectivity of directly photostimulated neurons (x-axis in panel j). Inset, difference in activity between photostimulation and non-photostimulation trials, averaged across all directly photostimulated neurons in a photostimulation group (Δ activity pg).**g.** Percentage of neurons with activity that is directly excited (gray) or coupled (black, excited; white, inhibited) by photostimulation, as a function of distance (from the nearest photostimulation target). **h.** Decay times (τ_pg_) of Δ activity pg (panel f, inset). Traces are color-coded based on decay time (gray lines). Green line, fluorescence decay after a brief burst of activity (Extended Data Fig. 7d). Magenta bar, time of photostimulation. **i.** Decay time constants of coupled neurons (τ_coupled_) vs. τ_pg_. **j.** τ_pg_ vs. average selectivity of the directly photostimulated neurons (panel f).

To probe the circuit basis of persistent activity we used two-photon photostimulation of small groups of neurons (Fig. 1b,c) and measured responses in other neurons (Fig. 1d-i). We targeted neurons in groups of eight (‘photostimulation group’), which ensured that we could 1) photostimulate sufficient numbers of spatially distributed neurons to potentially alter network activity and 2) observe changes in activity in non-targeted selective neurons, as an indicator of network connectivity. Targeted neurons were photostimulated by scanning the beam over their cell bodies for 3 milliseconds (Extended Data Fig. 4), causing short-latency (mean, 5 ± 2 ms; mean ± s.e.m.) spikes (range, 0.2 – 1.5 spikes per stimulus) (Extended Data Fig. 5). Neurons in photostimulation groups were photostimulated sequentially, ten times at 31.25 Hz (total duration, 319 ms; Extended Data Fig. 4). A large proportion of targeted neurons displayed increases in GCaMP6s fluorescence (ΔF/F; mean, 0.43; range, 0.07 – 0.80, 75% CI). Photostimuli were applied during the delay epoch (on 33.3 % or 40 % of trials). Multiple (2-5) photostimulation groups were stimulated during each behavioral session. Some groups contained mostly left-selective neurons (Fig. 1 d-f, top), whereas others were mainly right-selective (Fig. 1 d-f, bottom).

In addition to targeted neurons, cells up to 20 μm laterally from the center of the photostimulus could have been directly photostimulated (Fig. 1c, Extended Data Fig. 6). We refer to activated neurons in this neighborhood together as ‘directly photostimulated’ (Fig. 1f). Neural activity more than 30 μm from the targeted neurons changed as well (Fig. 1d). Control experiments show that changes in activity at distances greater than 30 μm from a photostimulus result exclusively from synaptic interactions with the directly photostimulated population (Fig. 1c, Extended Data Fig. 6). We refer to these neurons as ‘coupled’. Photostimulation groups produced detectable excitation in 20 coupled neurons (range, 4 – 39 neurons, 75% CI, two-tailed t-test, p<0.05), and inhibition in 6 coupled neurons (range, 0 – 13 neurons, 75% CI) per FOV. The number of excited or inhibited coupled neurons decreased with distance from the photostimuli (Fig. 1g) (length constant, 40 μm). Additional coupled neurons were presumably outside of the FOV.

Photostimuli caused transient increases in activity with diverse amplitudes and dynamics (Fig. 1e). On average, photostimulation increased the activity of targeted (Fig. 1e) and directly photostimulated (Fig. 1f) neurons for several seconds (mean, τ_pg_ = 3.1 s; range, 2.0 s — 4.6 s, 75% CI). The τ_pg_ are much longer than the decay of GCaMP6s fluorescence expected after a short burst of activity^12^ (τ_GCaMP6s_ = 1.1 s, Fig. 1h, green line; Extended Data Fig. 7). These long-lasting changes in fluorescence likely reflect persistent changes in spiking. Coupled neurons had decay times that were longer than τ_GCaMP6s_ and correlated with τ_pg_ (Fig. 1i, Pearson correlation = 0.49, p < 10^−5^), suggesting that photostimulation triggered self-sustaining activity. This activity was confined to a sparse subset of the persistently selective neurons. τ_pg_ were longest when the directly photostimulated population was selective for rightward licking (Fig. 1j, Pearson correlation = 0.28, p = 5×10^−5^). These data show that photostimulation of small groups of neurons can change activity in sparse populations of coupled neurons, and these changes outlast the photostimulus for several seconds.

Persistent activity is thought to be generated by recurrent connections between neurons with similar tuning^18,19^. We tested for specificity in functional connectivity by analyzing the responses of coupled neurons. We first analyzed photostimulated and coupled neurons based on their selectivity in trials without photostimulation. We determined whether the directly photostimulated population was mostly right-selective (R) (Fig. 2a, top) or left-selective (L) (Fig. 2a, bottom) and then analyzed (Fig. 2a, right) separately the responses of coupled R and L neurons. Responses were stronger in coupled R neurons when R neurons were photostimulated, consistent with like-to-like excitation (Fig. 2b, blue line; Pearson correlation = 0.15, p = 0.0025). However, coupled L neurons responded weakly to photostimulation of both R and L neurons (Fig. 2b, red line; p = 0.37). The like-to-like excitation from R neurons decreased steeply with distance (Fig. 2c, top), even though selective neurons were present throughout the field of view (Fig. 2c, bottom). These results suggest that recurrent excitation between sparse subsets of nearby R neurons (i.e. selective for contralateral movements) may contribute to maintenance of persistent activity. In addition, the strength of coupling was largest in neurons with trial-to-trial variability that was most correlated with the directly photostimulated neuronal population (Fig. 2d; Pearson correlation = 0.05, p < 10^−5^), suggesting that coupling reflected synaptic interactions, and consistent with like-to-like excitation.

**Figure 2:**
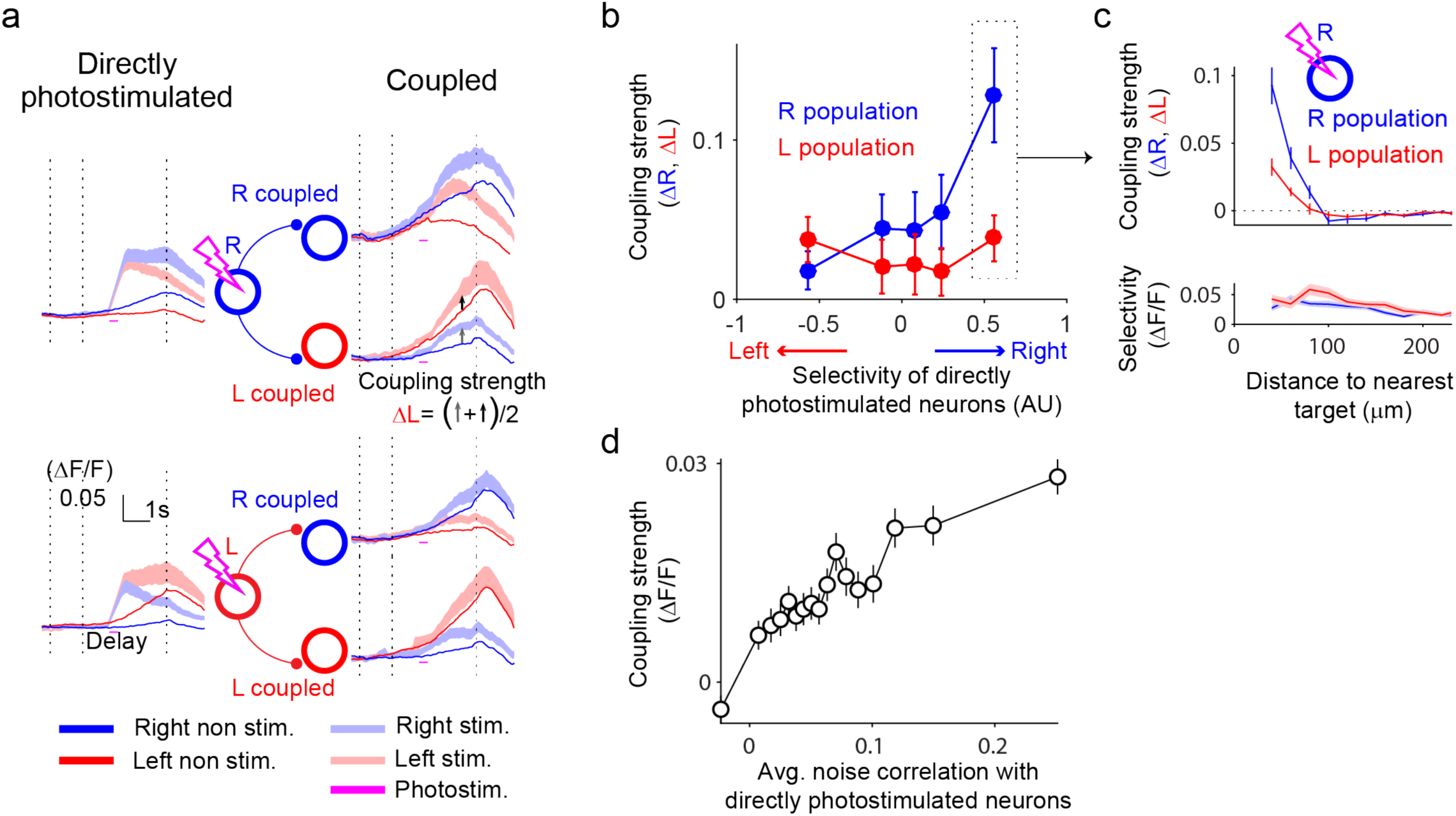
Functional connectivity depends on response type. **a.** Analysis of functional connectivity. Left, activity averaged across left-selective (L, red circle) and right-selective (R, blue circle) photostimulation groups (70 each; correct trials only). Dashed vertical lines denote the sample, delay and response epochs. Right, activity of coupled R or L populations. Traces, average activity of all coupled right-or left-selective neurons (30 — 100 µm from nearest target) weighted by the strength of their selectivity (S_i_^norm^, Methods). Arrows, calculation of coupling strength onto the L population (ΔL, Methods); calculation of ΔR is similar and based on R coupled neurons. **b.** Coupling strength (ΔR and ΔL) vs. average selectivity of directly photostimulated neurons (Fig. 1f). Data were binned in quintiles along x-axis (Error bars, s.e.m.). **c**. Top, Coupling strength (ΔR and ΔL) in response to photostimulation of R neurons vs. distance to nearest photostimulation target (corresponding to box in b.). Bottom, average selectivity of coupled R and L populations vs. distance to nearest photostimulation target. **d.** Coupling strength in each coupled neuron vs. its average correlation with all directly photostimulated

Selective delay epoch activity of ALM neural populations is causally linked to the direction of future licking^12^. Because targeted photostimuli produced changes in delay epoch activity that persisted until movement onset we tested for effects on animal behavior. A substantial proportion of photostimulation groups produced changes in behavior (p < 0.05, bootstrap; 22/215 and 35/215 photostimulation groups on right and left trials respectively) (Fig. 3a), nearly two-fold greater than expected by chance (Fig. 3b, Extended Data Fig. 8, p < 0.001 Kolmogorov-Smirnov test; Methods). These behavioral changes are consistent with other studies that have manipulated small numbers of neurons during behavior^20–22^.

**Figure 3:**
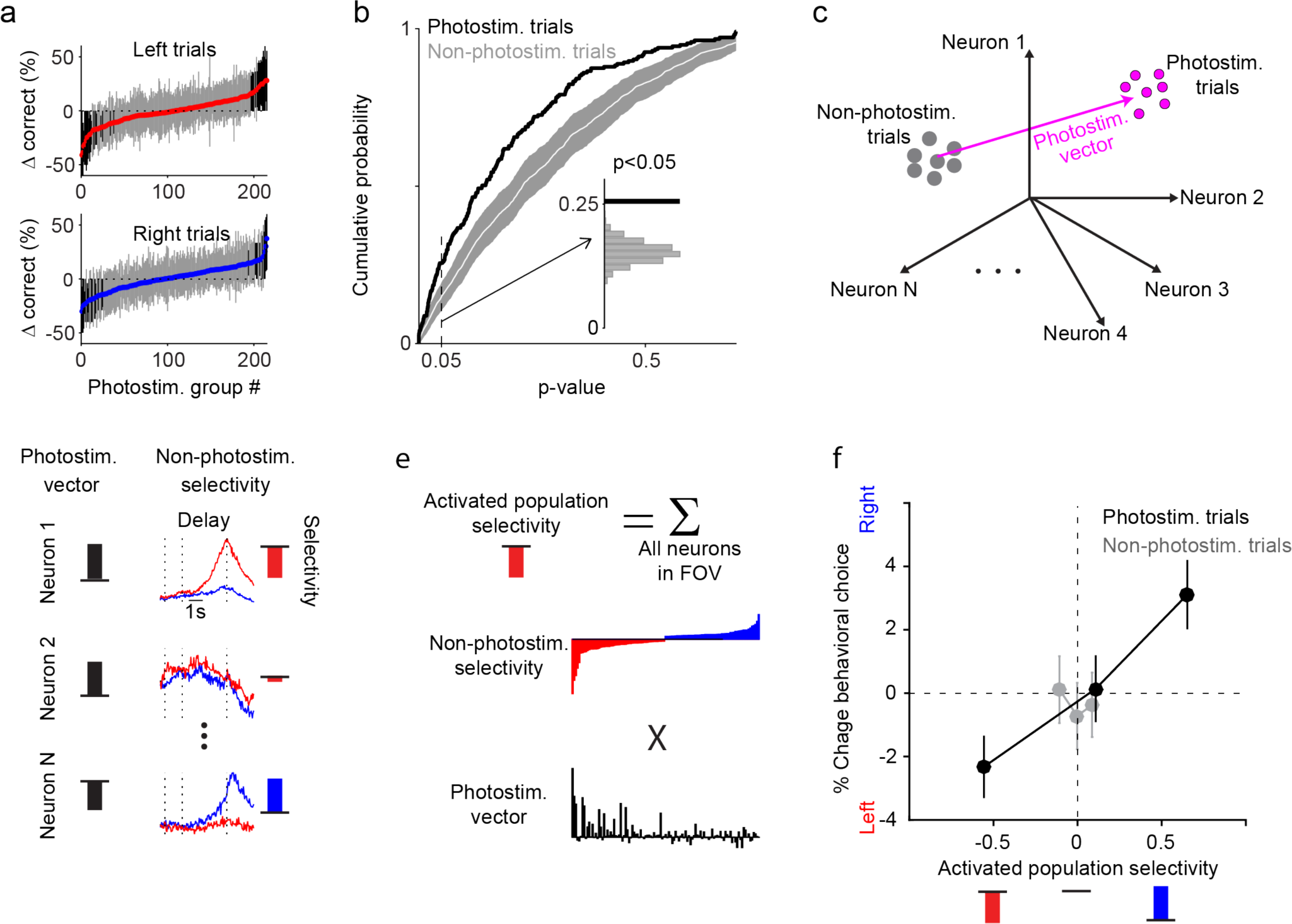
Photostimulation causes predictable changes in behavior. **a.** Change in behavioral performance between photostimulation and non-photostimulation trials for each photostimulation group for left (top) and right (bottom) trials. Bars represent 95% confidence intervals of the bootstrap. Black bars, p < 0.05. **b**. Cumulative distribution of p-values from individual sessions (bars from a) for photostimulation (black) and non-photostimulation (white) trials. Gray bar represents 95% confidence interval of bootstrap. Inset: Distribution of p < 0.05 for photostimulation (black) and non-photostimulation trials (gray). **c.** Schematic, photostimulated change in neural population in activity space. To take into account changes caused by photostimulation compared to trial-to-trial changes we computed the photostimulation vector (Methods). **d.** Left, photostimulation vector, contributions of individual neurons. Right, selectivity of individual neurons (S_i_, Methods). **e.** Activated population selectivity is the dot product of selectivity (S_i_) and the photostimulation vector. **f**. Change in percentage of right licking vs. activated population selectivity on photostimulation trials (Errorbars, s.e.m.).

To test how behavioral changes relate to selectivity of the photostimulation group, we calculated the net selectivity produced by the perturbation at the population level. For each imaged neuron we separately calculated the average photostimulated change in activity and the average selectivity. We then defined the ‘activated population selectivity’ as the product of these two quantities, summed across the population (Fig. 3c-e). Activated population selectivity was correlated with a bias in lick direction (Fig. 3f black line, Pearson correlation = 0.10, p < 0.01). These experiments show that targeted photostimulation of a small proportion of ALM neurons during the delay epoch has predictable effects on neural activity and behavior.

Measurements of low-dimensional neural activity in groups of neurons impose only weak constraints on network connectivity underlying neural computation^8^. Targeted photostimulation places additional constraints on circuit models. We found that common attractor models, monolithic line attractors^23^ and discrete attractors^24^, failed to capture key features of the responses to photostimulation: the sparse and persistent changes in activity (Fig. 4a, Extended Data Fig. 10).

**Figure 4:**
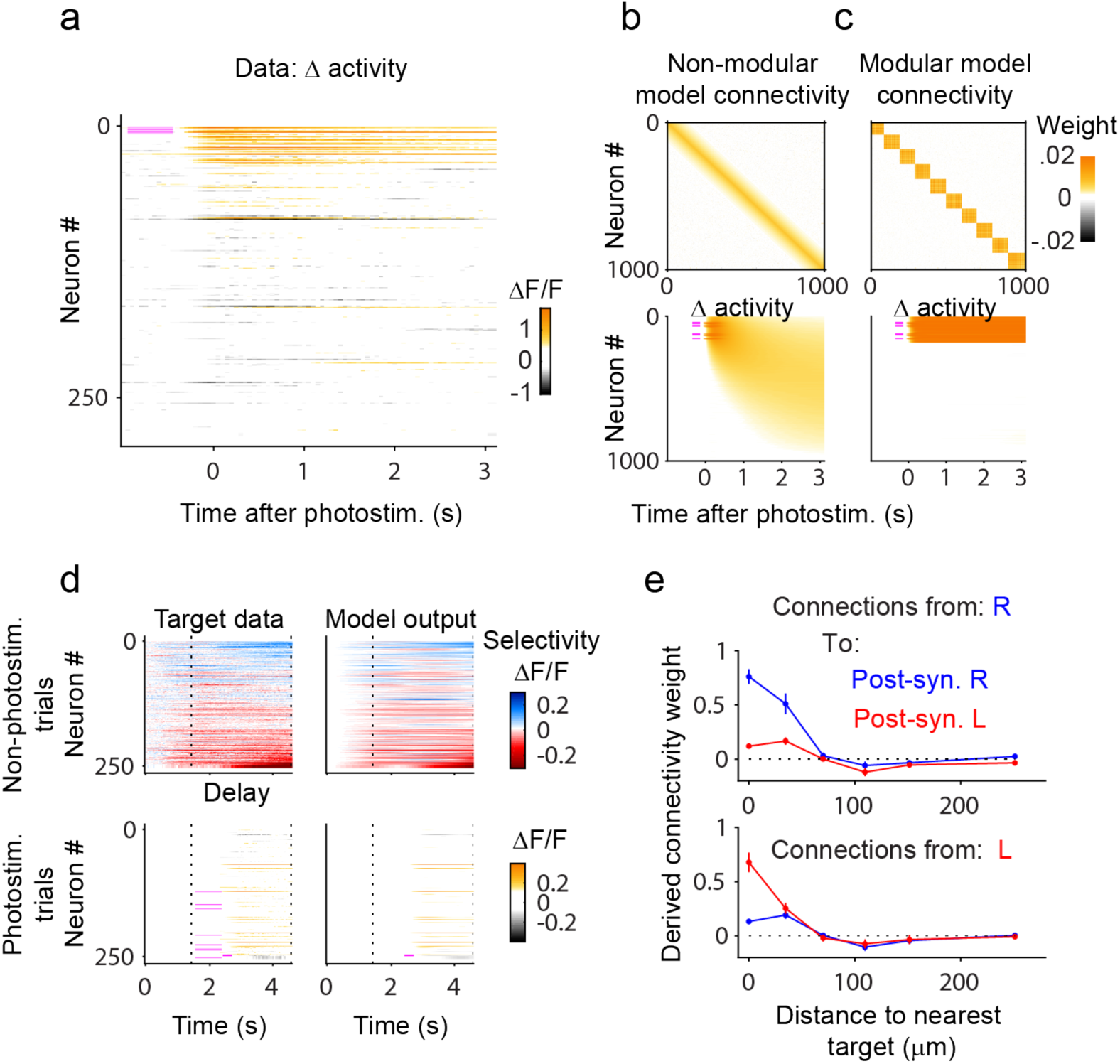
Modular network architecture explains sparse and persistent photostimulation responses. **a.** Δ activity of recorded neurons in one experimental session to one photostimulation group (8 neurons; magenta lines). **b-c.** Connectivity (top) and response to targeted photostimulation (bottom; magenta bars, photostimulated neurons) of model networks with locally biased connectivity with non-modular (**b**) and modular organization (**c**). Model neurons were sorted according to spatial location. **d**. Data-driven inference of connectivity. Model connections were tuned to match the model output (right panels) to the recorded data (left panels). Dashed lines denote the sample and delay epochs. **e**. Connection weights from presynaptic L (bottom panel) and R (top panel) neurons onto post-synaptic R (blue) and L (red) neurons as a function of distance between neurons.

Imposing sparse, local connectivity on such models did not produce sparse, persistent activity (Fig. 4b). We tested a ‘modular’ network, with strong within-module connectivity and weak between-module connectivity (Fig. 4c), in which each module can independently maintain persistent activity. Such a network as a whole produced sparse persistent activity consistent with experiments.

To further test if this modular architecture is consistent with targeted photostimulation experiments, we fit^25^ model networks directly to the population recordings in photostimulated and non-photostimulated trials (Fig. 4d). The resultant inferred networks were biased towards local connectivity (Fig. 4e), reflecting the length scale of the photostimulation-response (Fig. 1g, Fig. 2c) and local cortical connectivity^26,27^. Connections between R neurons were stronger than connections between L neurons (Fig. 4e) enabling the inferred networks to produce longer time scale responses to photostimulation of R neurons than L neurons (Fig. 1j). Analysis of the local connectivity revealed that the inferred networks resemble the modular network in Figure 4c (Extended data Fig. 10). Networks trained to match only the non-photostimulation trials exhibited significantly weaker connections that resembled the monolithic network (Fig. 4b), demonstrating the power of perturbation experiments to constrain neural network models.

We used calcium imaging simultaneously with targeted photostimulation to probe coupling of functionally identified groups of neurons during behavior. These experiments revealed “like-to-like” connectivity between R neurons in the left hemisphere of ALM, and less tuned excitation for L neurons, consistent with previous experiments showing stronger noise correlations in spiking between R neurons than L neurons^15^. Transient perturbations of small groups of neurons caused network responses lasting several seconds, modifying the internal state of a key-circuit for decision-making and modulating behavior seconds later. Even when considering neurons with similar functional properties in a neighborhood, the evoked persistent responses were limited to a subset of neurons. In standard attractor models, activation would spread globally. We propose modular networks with dynamics that are consistent with the data. We hypothesize that these networks may be less sensitive to noise than line-attractor networks, and have higher memory capacity than discrete attractor networks.

Our finding that local photostimulation produces persistent changes can be reconciled with previous studies showing that ALM dynamics recover rapidly from network-wide photoinhibition^18,19^. For example, the shapes of within-module energy troughs could have a flat bottom; strong, network-wide perturbations could selectively engage inter-modular dynamics or non-linear interactions with different circuits (e.g. thalamocortical interactions^28^; Extended data Fig. 10). The wide range of length scales over which interactions produce robust, attractor-like dynamics in our models motivate future experiments in which larger numbers of neurons can be photostimulated^29,30^ over varying length scales.

## Acknowledgements

We thank R. Darshan, B. Mohar, A. Finkelstein, H. Inagaki, S. Romani, A. Singh, M. Pachitariu and S. Peron for comments on the manuscript; T. Pluntke and R. Mohar for animal training; X. Zhang, K. Ritola and H. Inagaki for making the ST-chrimsonR constructs; E. Fardone for histology; and P. Rickgauer for discussions. This work was funded by Howard Hughes Medical Institute. KD is a Helen Hay Whitney Foundation postdoctoral fellow and was supported by the Simons Collaboration on the Global Brain.

## Extended Data Figures

**Extended Data Figure 1:**
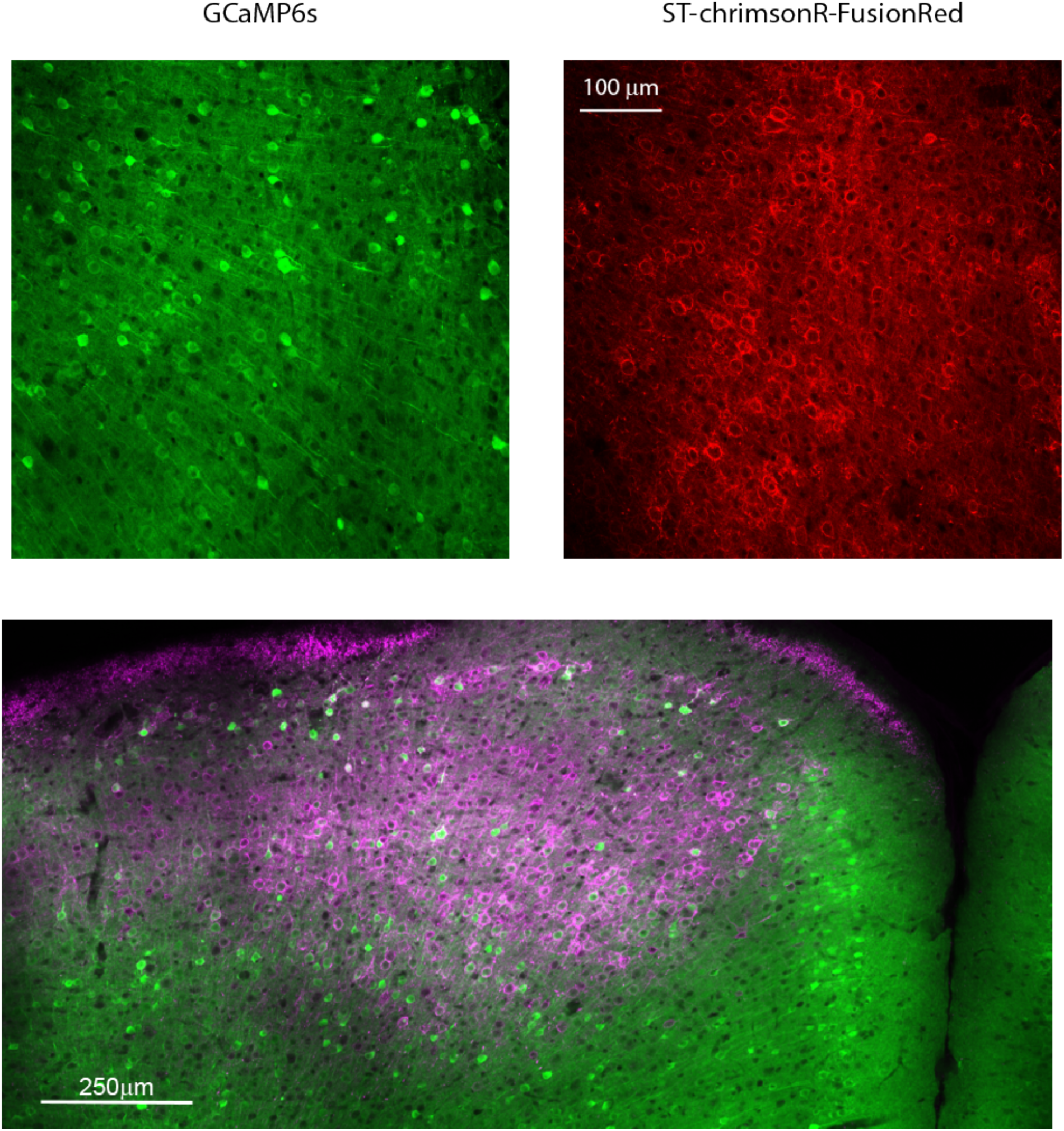
Coexpression of GCaMP6s and ST-chrimsonR. After experiments brains were harvested and sectioned (100 µm coronal sections). Images show co-expression of GCaMP6s (green) and ST-chrimsonR-FusionRed (red) from a section that was under the cranial window.

**Extended Data Figure 2:**
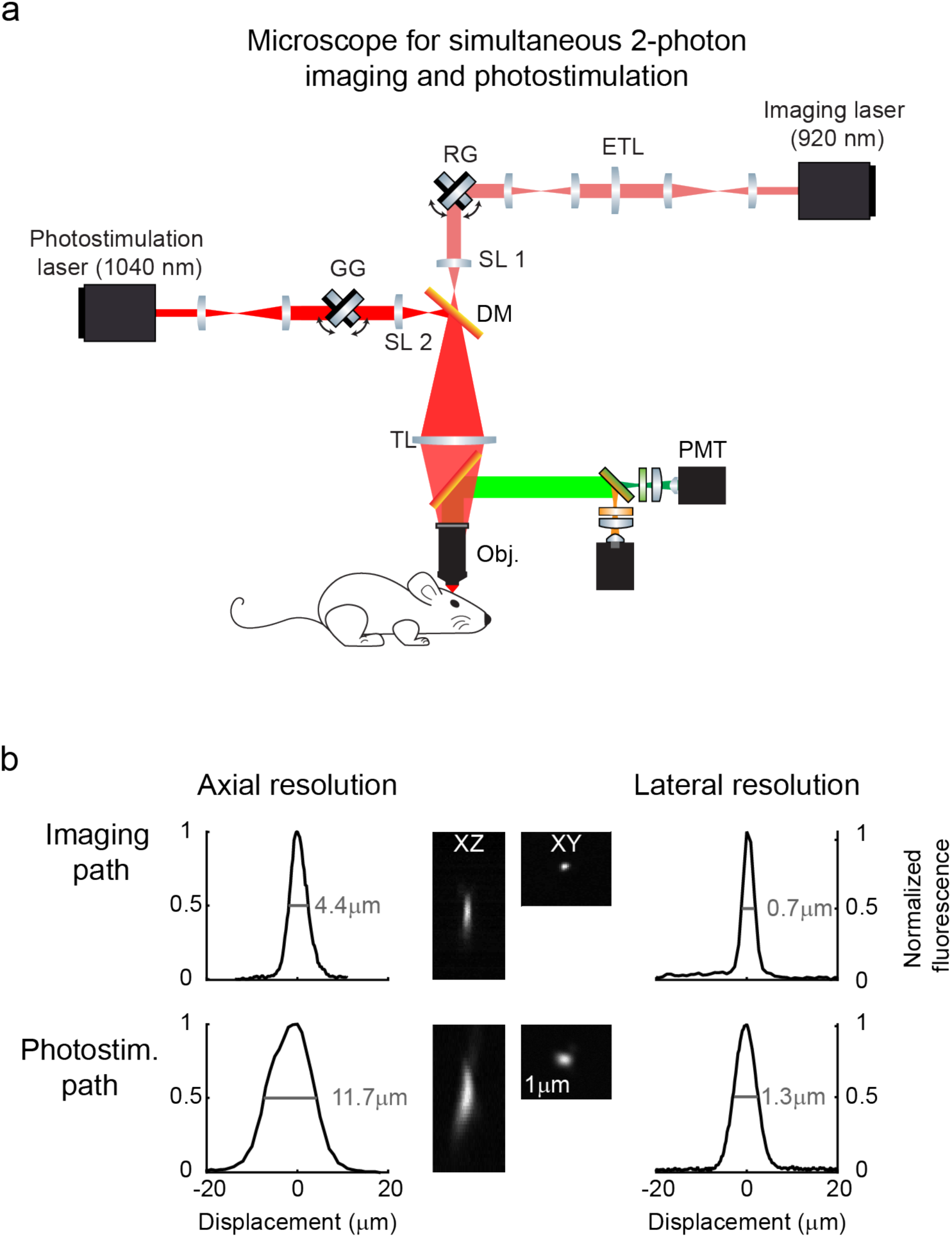
Simultaneous imaging and targeted photostimulation. **a**. Schematic of the microscope: Photostimulation laser, 1040 nm (Fidelity HP, Coherent); Imaging laser, 920 nm (Chameleon Ultra II, Coherent); GG, pair of 3 mm galvanometer mirrors (Cambridge, 6215H); ETL, electric tunable lens, (EL-10-30-C, Optotune); SL 2, scan lens photostimulation path, 33 mm focal length, a stack of 3 x 100 mm focal length lenses (AC-254-100b, Thorlabs); SL 1, scan lens imaging path, 30 mm focal length (55-S30-16T, Special Optics); DM, 1000 nm short-pass dichroic mirror (Edmund optics); TL, Tube lens 160 mm focal length (Special Optics); Obj., 16x objective, 0.8 NA, 3mm working distance (CFI75 LWD, Nikon); PMT, photomultiplier tubes (H10770(P)B-40, Hamamatsu). **b**. Point-spread functions. Measurements were made by imaging 500 nm yellow-green beads (Polysciences). Reported values correspond to the full-width at half maximum.

**Extended Data Figure 3:**
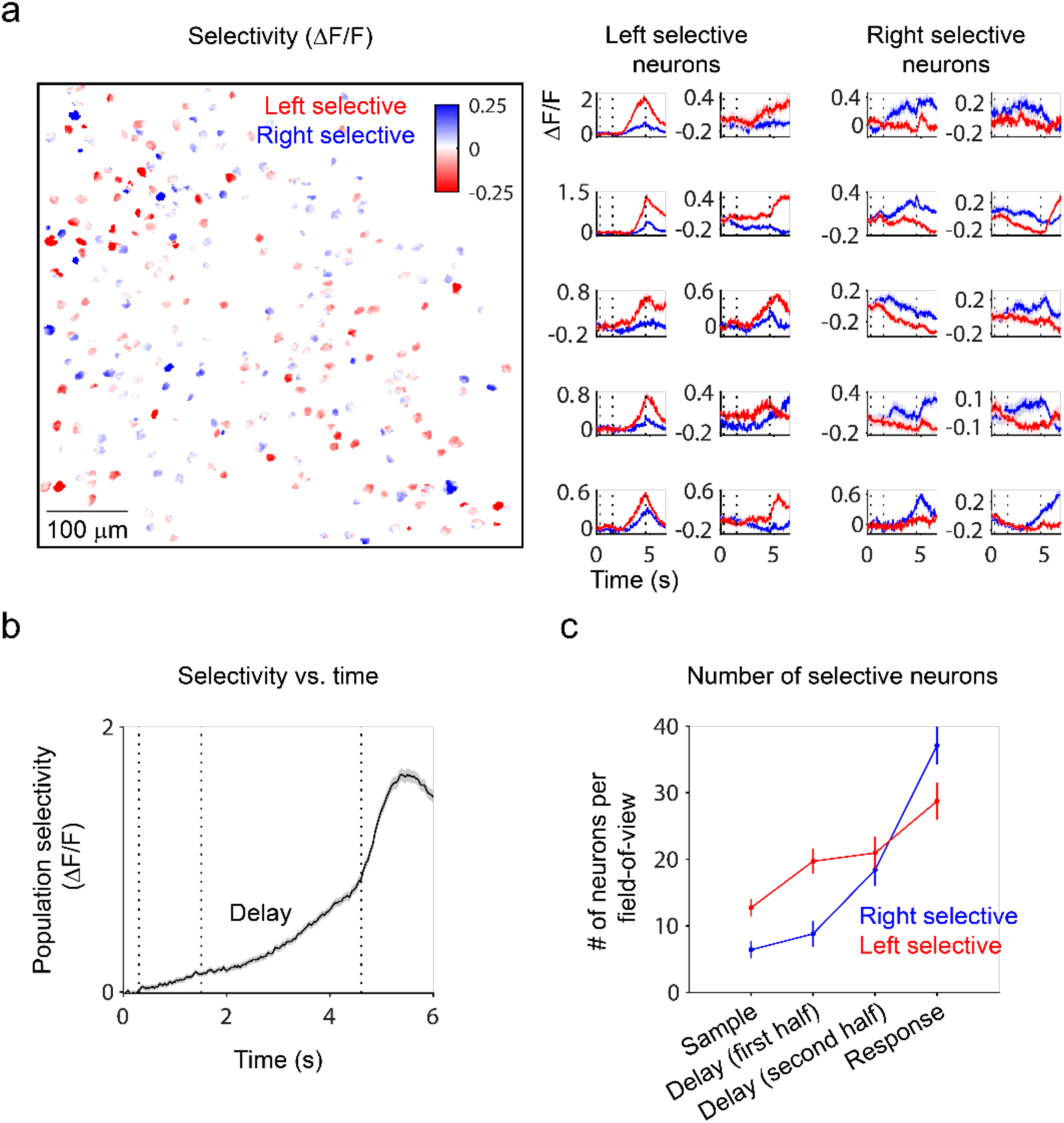
Trial-type selectivity. **a.** Map of selectivity for individual neurons in one behavioral session. Selectivity was calculated as the difference in activity on correct right and left trials at the end of the delay epoch. Selective neurons were scattered across the field of view (left panel), and displayed heterogenous dynamics (right panel, example neurons). Dashed vertical lines denote the sample, delay and response epochs. **b.** For each behavioral session all neurons were categorized based on their selectivity at each time point:

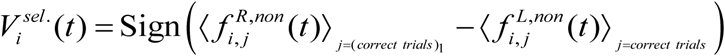

where trial averaging was performed over a randomly-chosen training subset of 20 % of correct trials denoted as “correct trials_1_”. With the remaining 80% of trials (correct trials_2_, testing subset) we computed the population selectivity as:

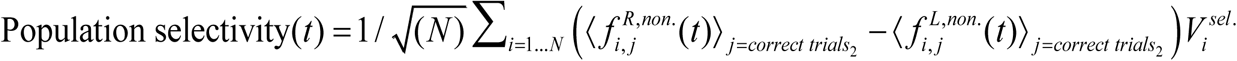

where N is the number of neurons in the FOV. Dashed vertical lines indicated the sample, delay and response epochs, Errorshade is the s.e.m. across sessions. **c.** We used a T-test to compare the epoch-averaged activity on left and right trials for each neuron:

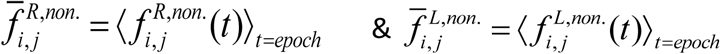 We then counted the number of neurons for which the resulting p-value was less than 0.05.

**Extended Data Figure 4:**
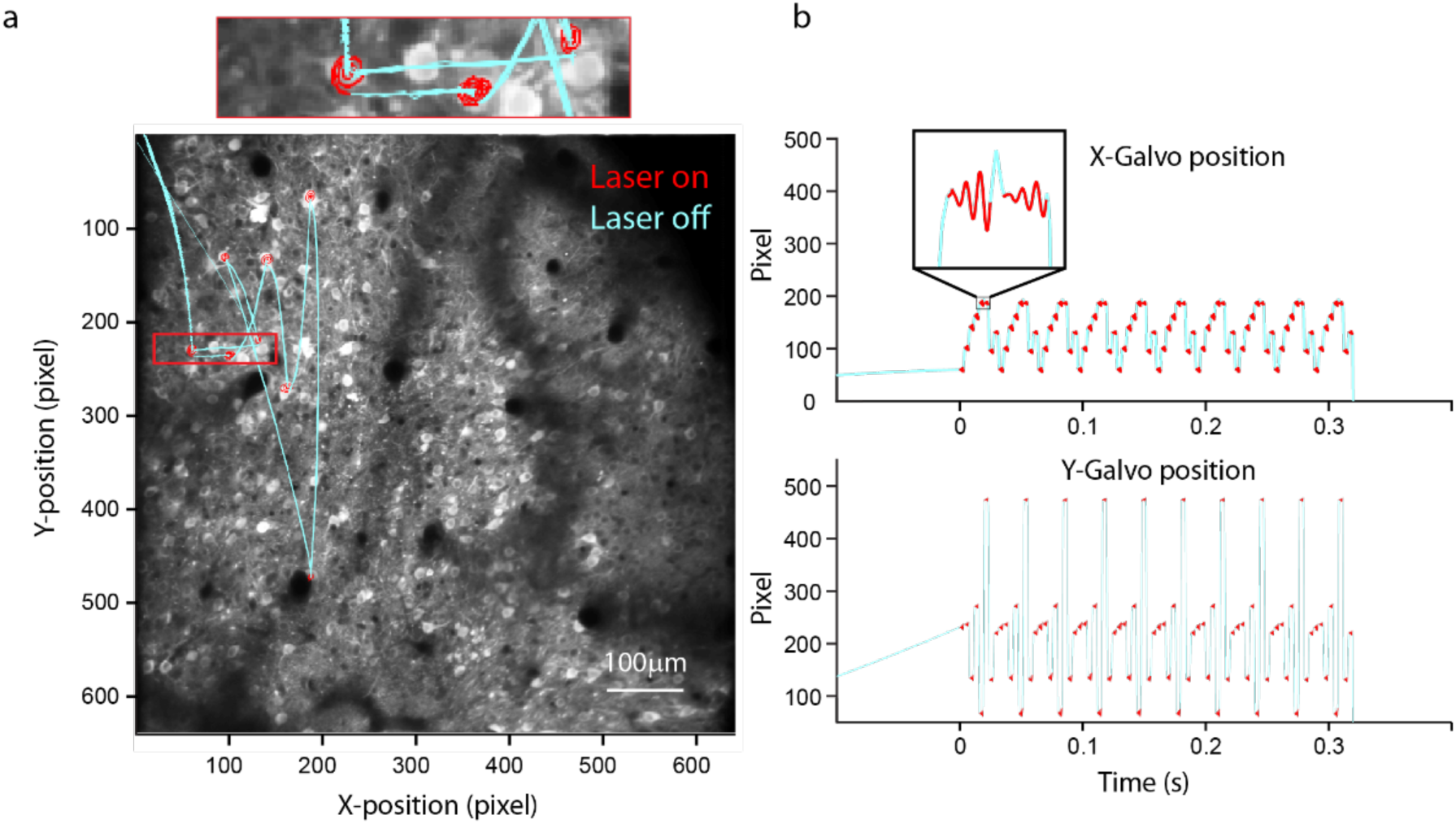
Photostimulation scan trajectory. **a**. Photostimulation was performed by scanning the laser over a target neuron, defined by GCaMP6s fluorescence, in a spiral pattern. The image shows the scan path, reconstructed from position encoders of the galvo mirrors, superposed on a fluorescence image. Each spiral (red) lasted for 3 ms. Laser power was then turned off for 1 ms while the beam was redirected to the next neuron in the sequence (cyan). Once the last (8^th^) neuron was photostimulated the beam was directed back to the first neuron. This sequence was repeated 10 times per photostimulation trial (**b**).

**Extended Data Figure 5:**
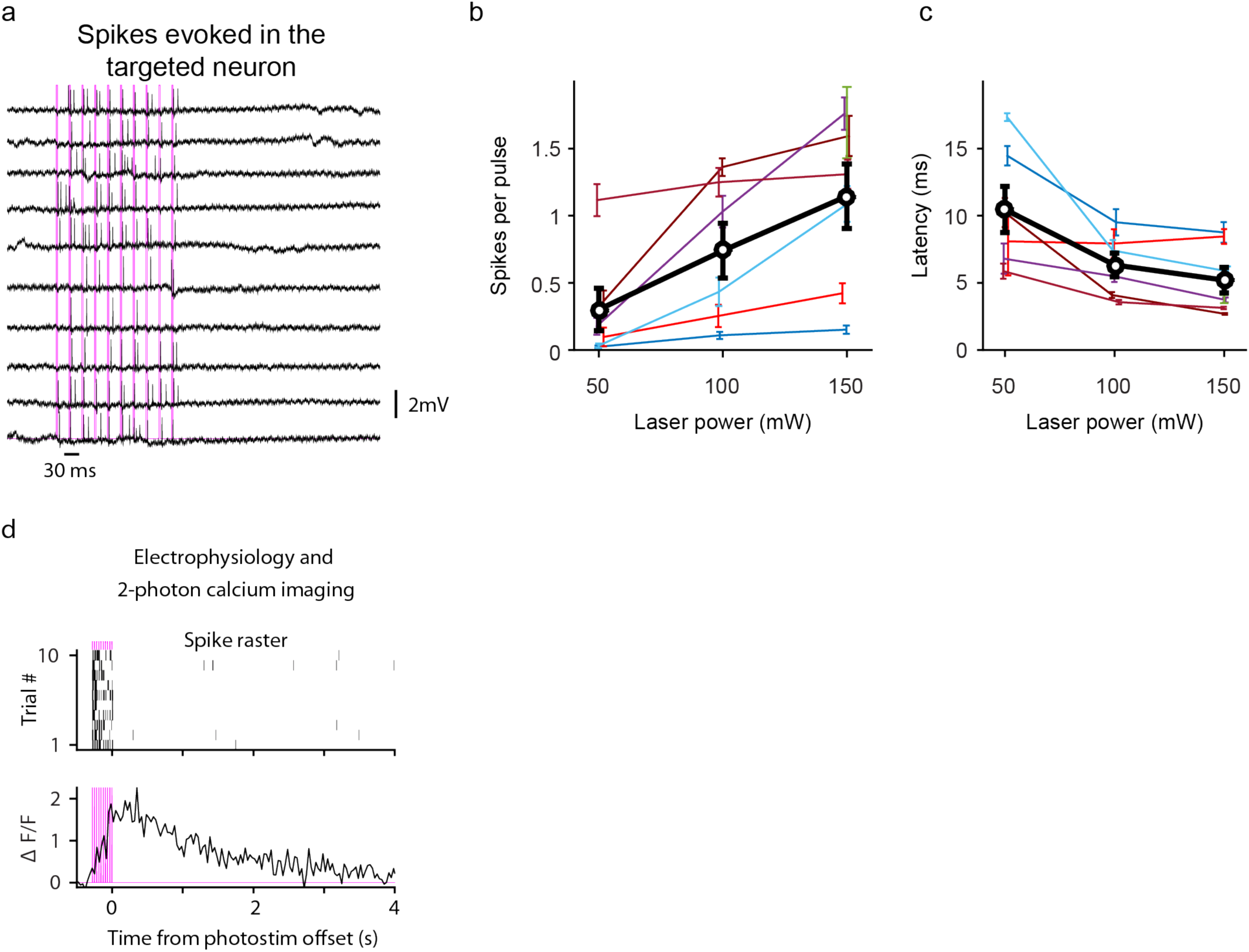
Electrophysiology during targeted photostimulation. Loose-seal, cell-attached recordings during photostimulation in anesthetized mice. Single neurons (n=7) were photostimulated using the pattern shown in Extended Data Figure 4a with either 50, 100 or 150 mW laser power. **a**. Photostimuli produced increases in spiking. Red bars, photostimuli. **b**. Spikes evoked by single photostimuli (spiral) as a function of laser power. The weakest photostimuli (50 mW) failed to drive spikes in most neurons. All photostimulation experiments done during behavior used either 100 or 150 mW, which evoked on the order of one spike per photostimulus. **c**. Spike latency as a function of laser power. The spike latency for 150 mW was 5±2 ms (mean ± s.e.m.). **d**. Simultanous GCaMP6s fluorescence and extracellular voltage, showing that photostimulation produced increases in spiking that were associated with large increases in fluorescence. These fluorescence transients decayed with a 1 s time constant, consistent with previous observations^1^, and much faster than decays observed when photostimulating groups of neurons during the delay epoch.

**Extended Data Figure 6:**
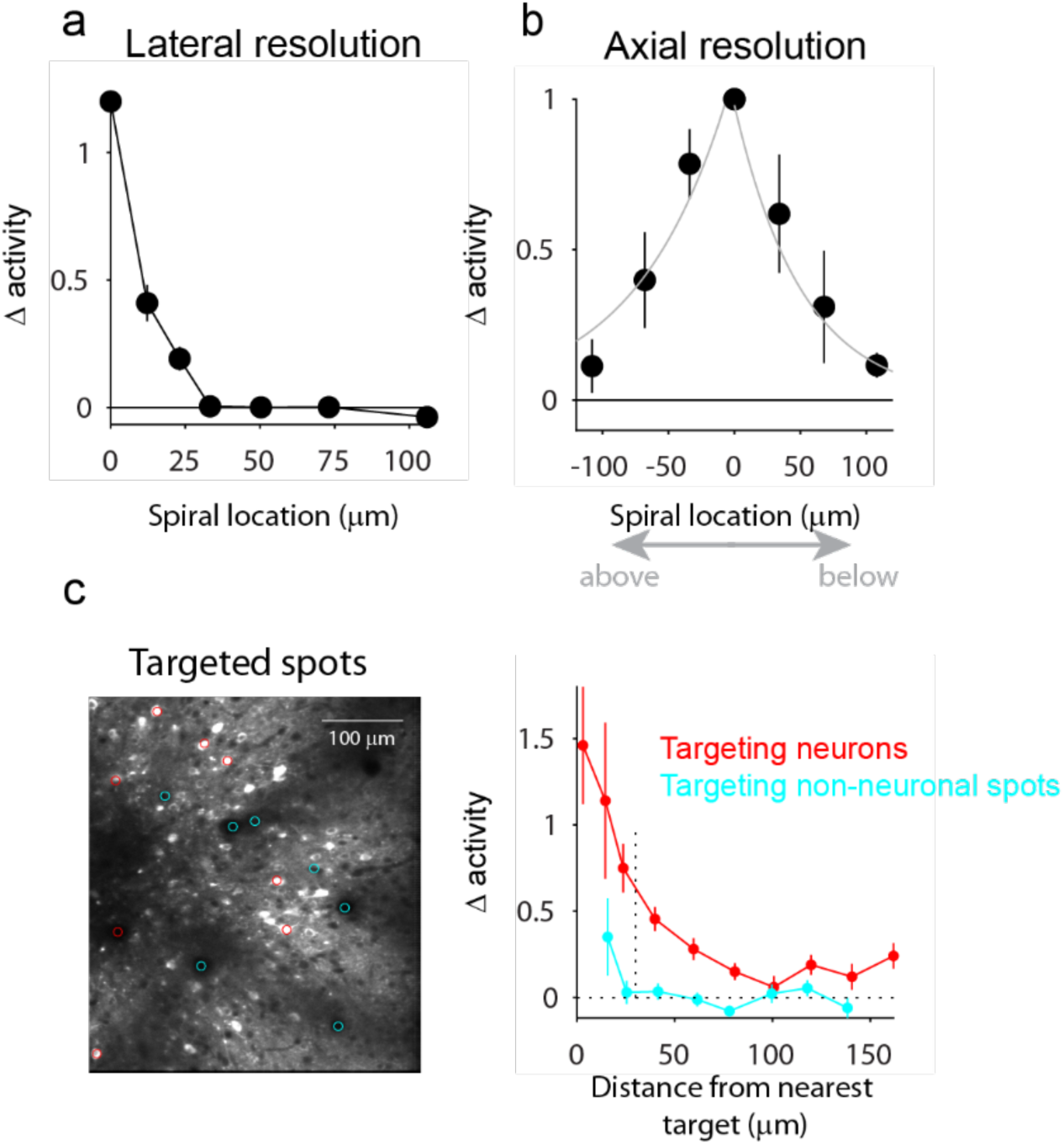
Spatial resolution of photostimulation. **a**. Lateral resolution. Resolution was characterized during quiet wakefulness by measuring the amplitude of photostimulation-induced GCaMP6 transients with the photostimulus at varying distances from the recorded neuron (n = 9 neurons, 10 trials for each photostimulus location). The normalized change in activity was calculated as:

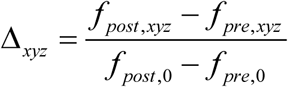

where *f*_*pre,xy*_ is the average fluorescence before photostimulation (150 ms) and *f*_*post, xy*_ is the fluorescence averaged immediately after the photostimulation (1 s) at a lateral distance of xy micrometers from the imaged cell. Changes in activity were limited to a radius of less than 30 micrometers. **b**. The axial resolution in these experiments is 80 micrometers (FWHM), about a factor of 1.5 or 2 larger than reported using similar methods^2,3^. This broadening is because we used stronger photostimuli to produce reliable changes in activity in the targeted neurons. Given variable levels of expression and excitability, these laser powers were sufficient to drive spikes even in weakly expressing neurons, at the cost of degraded axial resolution^4^. **c.** Additional resolution measurements in awake mice. Resolution experiments described above were performed during periods of quiet wakefulness. To further account for potential differences in neuronal excitability during behavior we performed additional experiments to estimate the lateral spatial resolution of group photostimulation during behavior. As in Figure 1, we selected groups of 8 neurons to photostimulate during behavior (red). Additionally, we selected groups of 8 non-neuronal spots to target for photostimulation (cyan). For the groups of non-neuronal spots, we find that there is weak activation of neurons within 20 microns of a target, but no significant changes were observed more than 30 microns away from a target location. Because photostimulation of non-neuronal spots fails to produce changes in activity at distances greater than 30 microns from the photostimulation target, we treat any neurons located 30 microns or more from a target throughout the text as “coupled”. In addition, we treat any neurons located within 20 microns of a target whose activity was significantly altered as “directly photostimulated”.

**Extended Data Figure 7:**
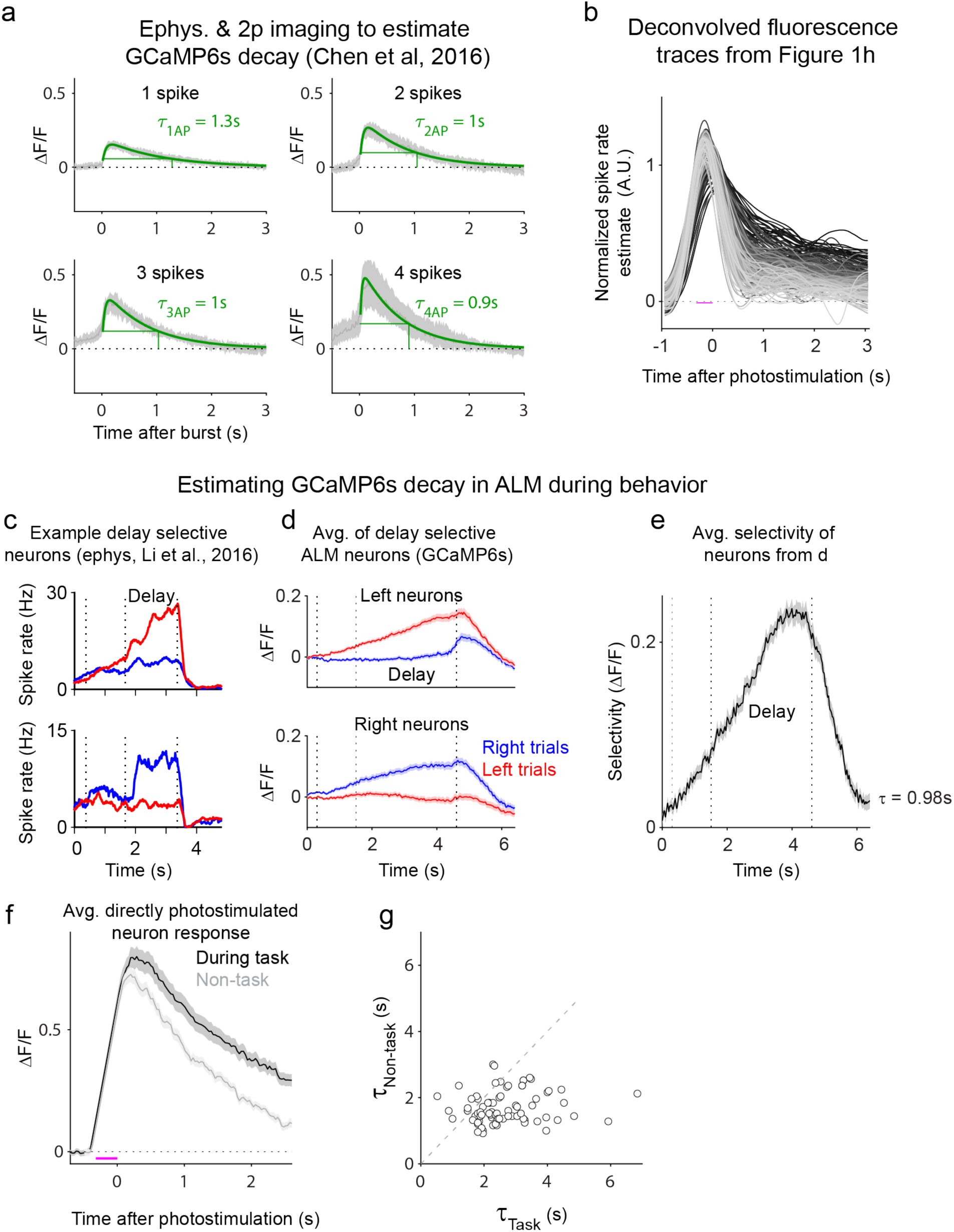
Dynamics of activity-dependent fluorescence. **a**. Estimating GCaMP6s decay after short bursts of activity in anesthetized mice. We analyzed simultaneously recorded cell-attached electrophysiology and two-photon imaging of GCaMP6s in the visual cortex^1,5^ to assess the decay of GCaMP6s fluorescence. The decay time constant is the time at which the activity decays to 1/e of its peak value. We analyzed the fluorescence traces following bursts of 1, 2, 3 or 4 action potentials and found τ’s of 1.2s, 1s, 1s and 0.9 s respectively. **b.** We fit each fluorescence response with the kernel *Κ*_*GCaMP*6*s*_ (green traces) according to the equation:

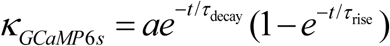

and obtained the following average parameter values: a = 0.31 τ_decay_ = 0.87 s and τ_rise_ = 0.06 s. We used *Κ*_*GCaMP*6*s*_ to estimate the spiking dynamics underlying the fluorescence responses of directly photostimulated neurons by deconvolving each trace in Figure 1h with the kernel *Κ*_*GCaMP*6*s*_. Prior to deconvolution, traces were smoothed with a cubic smoothing spline (MATLAB; csaps, P = 0.96). We found that 95% of photostimulation groups produce changes in estimated spike rate that remain elevated 2 seconds or longer following photostimulation. **c.** We also used known rapid spike rate changes in ALM to estimate the GCaMP6s decay time during behavior^6^. ALM contains neurons in which selectivity peaks at the end of the delay epoch and rapidly decays to zero following the response cue. Example neurons with late-delay right (top) and left (bottom) selectivity^7^ are shown. Fluorescence dynamics in these neurons will enable us to estimate the GCaMP6s decay time constant by observing the decay of selectivity following the go-cue. **d.** We identified neurons with late delay epoch selectivity (Two-tailed T-test, p<0.2) that was greater than the response response epoch selectivity (Two-tailed T-test, p>0.2). On average we found 13.5 such neurons per session (6 — 22). The traces show the average of right (top) and left (bottom) selective neurons. **e**. We next plotted the average selectivity of these delay selective neurons as calculated in Extended Data Fig. 3. From the electrophysiological recordings, we assume that spike rate selectivity of these neurons drops to zero. We found that selective fluorescence decayed with τ = 0.98 s, consistent with the measured decay times of GCaMP6s in **a**. This is expected to be an upper bound on the GCaMP6s fluorescence decay time because selectivity in some of these neurons may not instantaneously drop to zero and produce erroneously long estimates of τ. **f**. To test if the persistent changes in activity could also be evoked during periods of quiet wakefulness, on a subset of experiments we photostimulated the same groups of neurons immediately after the animal completed the behavioral session (3 mice, 10 photostimulation groups). We found that the activity averaged across all groups and directly photostimulated neurons decayed more slowly during the task than in the non-task period. **g.** For each directly photostimulated neuron (84 neurons), we calculated the decay time constant both during the task and during the non-task period. Decay time constants during the task (τ = 3.1 s, 1.8 s — 4.3 s, 75% CI) were significantly longer than time constants outside of the task (τ = 1.7 s, 1.2 — 2.4, 75% CI; p < 10^−5^, two-tailed T-test; errorbars, s.e.m.).

**Extended Data Figure 8:**
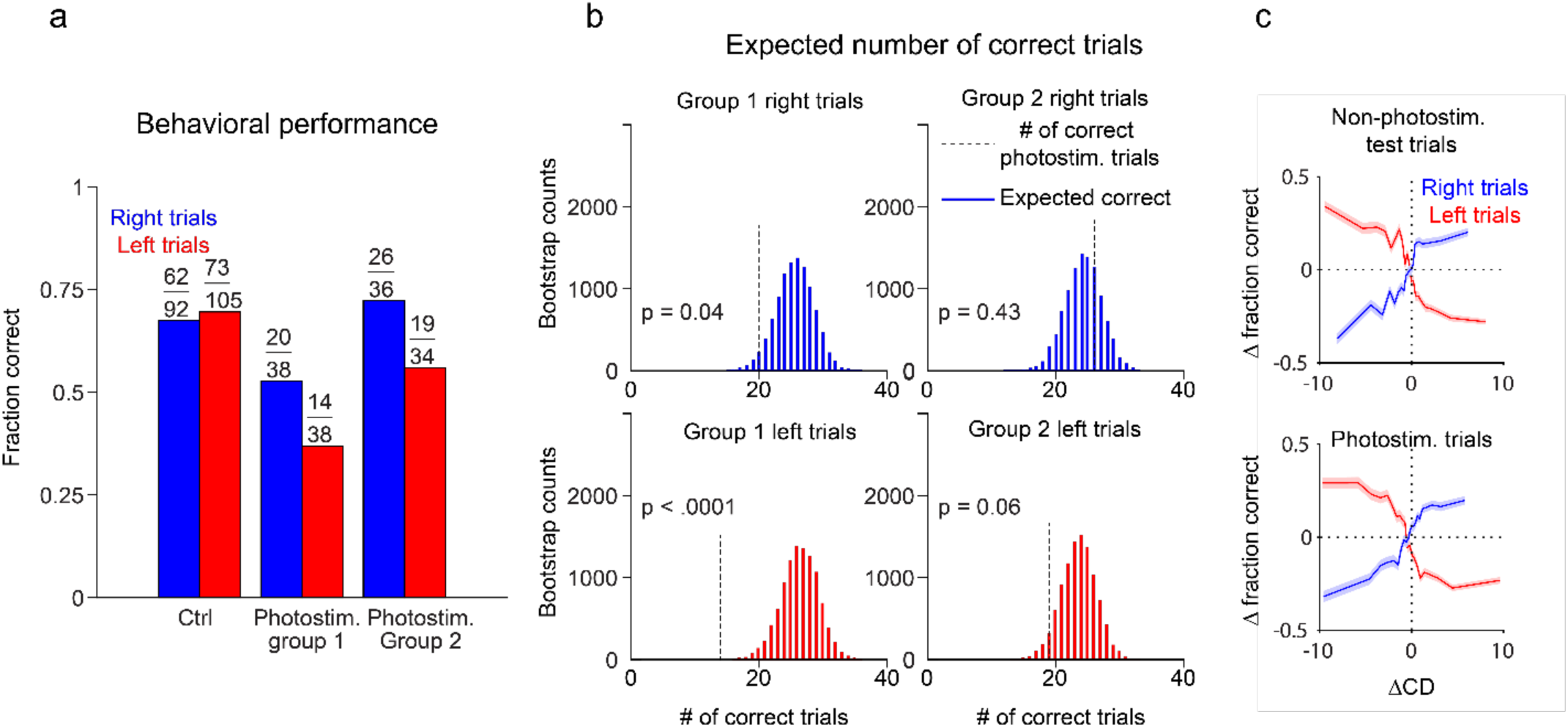
Effect of photostimulation on behavior. **a.** Behavioral performance of a mouse on a single session. Performance dropped from 73/105 correct (69%) on non-photostimulation lick left trials to 14/38 (37%) when photostimulation group 1 was photostimulated. **b.** A bootstrap distribution of performance on non-photostimulation trials was generated by randomly sampling 38 of the 105 non-photostimulation trials 10,000 times (b, bottom left). For each of the 10,000 random samplings, we counted the number of correct trials to determine the p-value for each photostimulation group. Bootstrap distributions were used to determine the 95% confidence intervals in Figure 3a. **c.** To relate changes in performance to changes in activity, we projected population activity along a “coding direction” (CD) which maximally separates trial-averaged activity on a subset of “training” left and right non-photostimulation trials^8^. We then projected single-trial activity from a testing subset of non-photostimulation trials (top) as well as photostimulation trials (bottom) along the CD and calculate the average change in performance vs. displacement along the CD. The relationship between CD activity and behavior was similar for photostimulation and non-photostimulation trials because photostimulation triggered only sparse changes in activity.

**Extended Data Figure 9.**
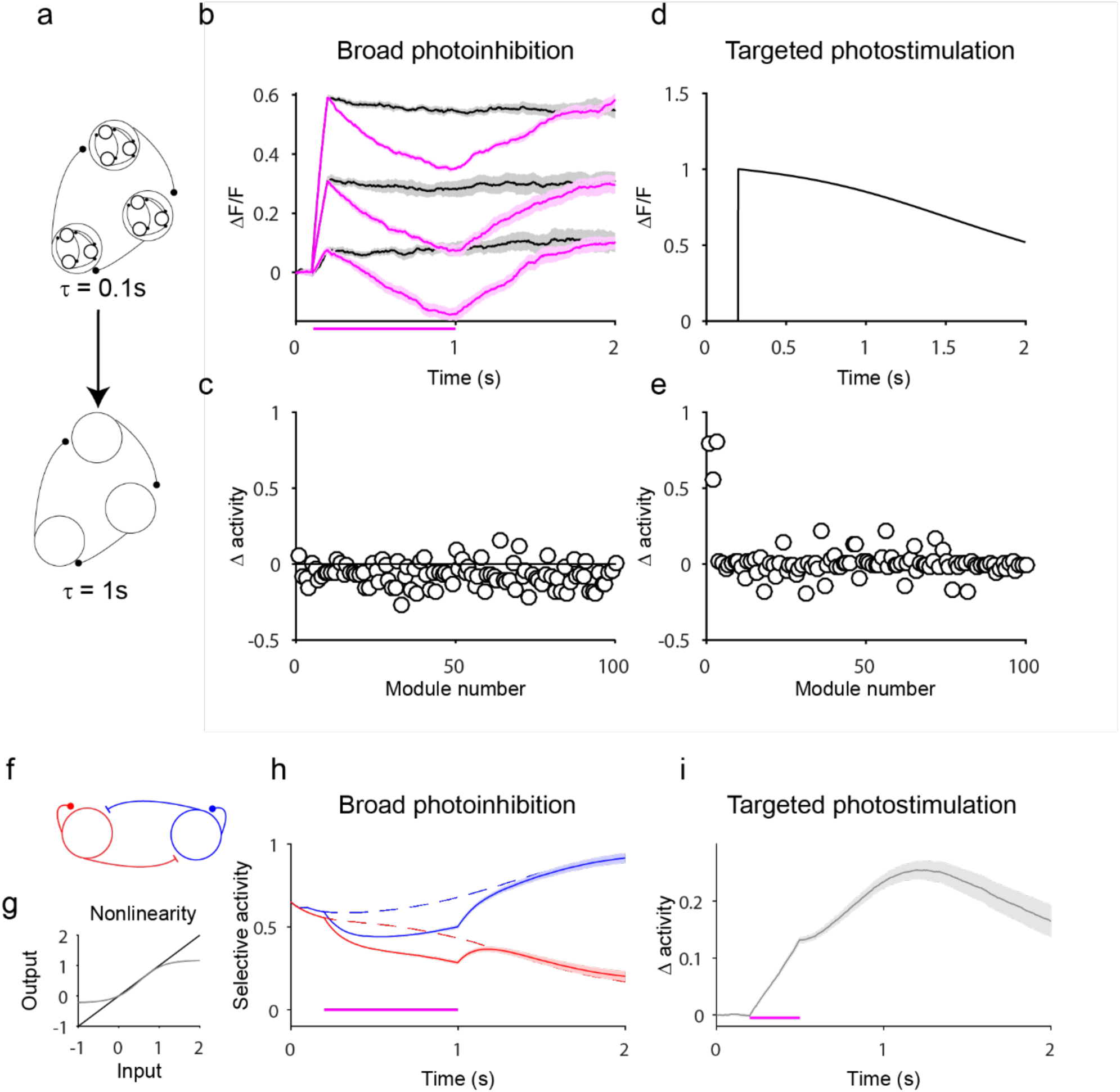
Network models of robustness. The model in Figure 4d-e produces persistent selectivity and sparse persistent responses to photostimulation through a locally connected modular architecture. The model was fit directly to the data, and all of the relevant connections for maintaining persistence are between a small number of neighboring neurons. The toy model network in Figure 4c illustrates this modularity. This model produces sparse persistent responses to photostimulation of a small number of neurons. However, this model does not by itself produce the rapid recovery from broad photoinhibition that has been described in ALM^9^. One possible reconciliation of these two findings can be achieved by tuning intermodular connections to produce rapid recovery from broad photoinhibition. **a.** We made a simplified network model in which individual modules are represented as units with 1 second synaptic time constants. These long time constants reflect the strong within module feedback. The spike rate of module *i* was given by the equation:

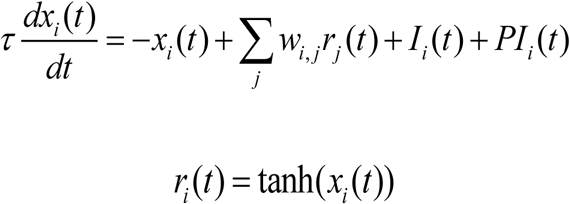 Where *τ*= 1 s, and the external input *I* was equal to 1 during the sample period and 0 for all other time points. The network was trained using the FORCE algorithm^10^ to produce a persistent output along one dimension. On alternating training trials, broad transient photoinhibition given by PI(t) was applied immediately after the sample period. Training sought to produce a change in activity along the persistent mode during photoinhibition followed by a rapid recovery after the offset of photoinhibition. **b.** We found that the trained network not only matched the target output, but also produced graded persistent activity in response to varying the amplitude of the external inputs. Magenta traces show the response of the network to broad photoinhibition illustrating the network’s robustness to these particular perturbations. **c.** To illustrate the widespread nature of the photoinhibition, we plotted the change in activity resulting from photoinhibition for each module. **d.** Finally, because of the long intrinsic modular time constants, we found that targeted photostimulation of small numbers of modules produced changes that persisted for several seconds. **e**. To assess whether targeted photostimulation cause sparse changes in network activity as in our data (Fig. 1g, Fig. 4a) we plotted the change in activity due to photostimulation for each module. The amplitude of excitation in the three targeted modules is significantly larger than in all other modules. **f.** A hybrid discrete/continuous attractor model^8^, can also produce rapid recovery to broad photoinhibition and persistent changes from targeted photostimulation. In this model, a fined-tuned approximately linear transfer function 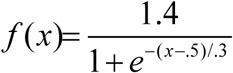 (**g**) coupled with self-exciting and mutually inhibiting L (red) and R (blue) neurons produces slow dynamics along the difference mode [1, −1] and fast dynamics along the [1, 1] direction. Large amplitude inhibition of both neurons produces a rapid recovery (**h**) whereas weak excitation of one of the neurons produces a persistent change in activity (**i**). Because this network is only composed of 2 units, it does not produce sparse and persistent responses, but a network composed of many such modules would produce sparse responses.

**Extended Data Figure 10. Monolithic models.**
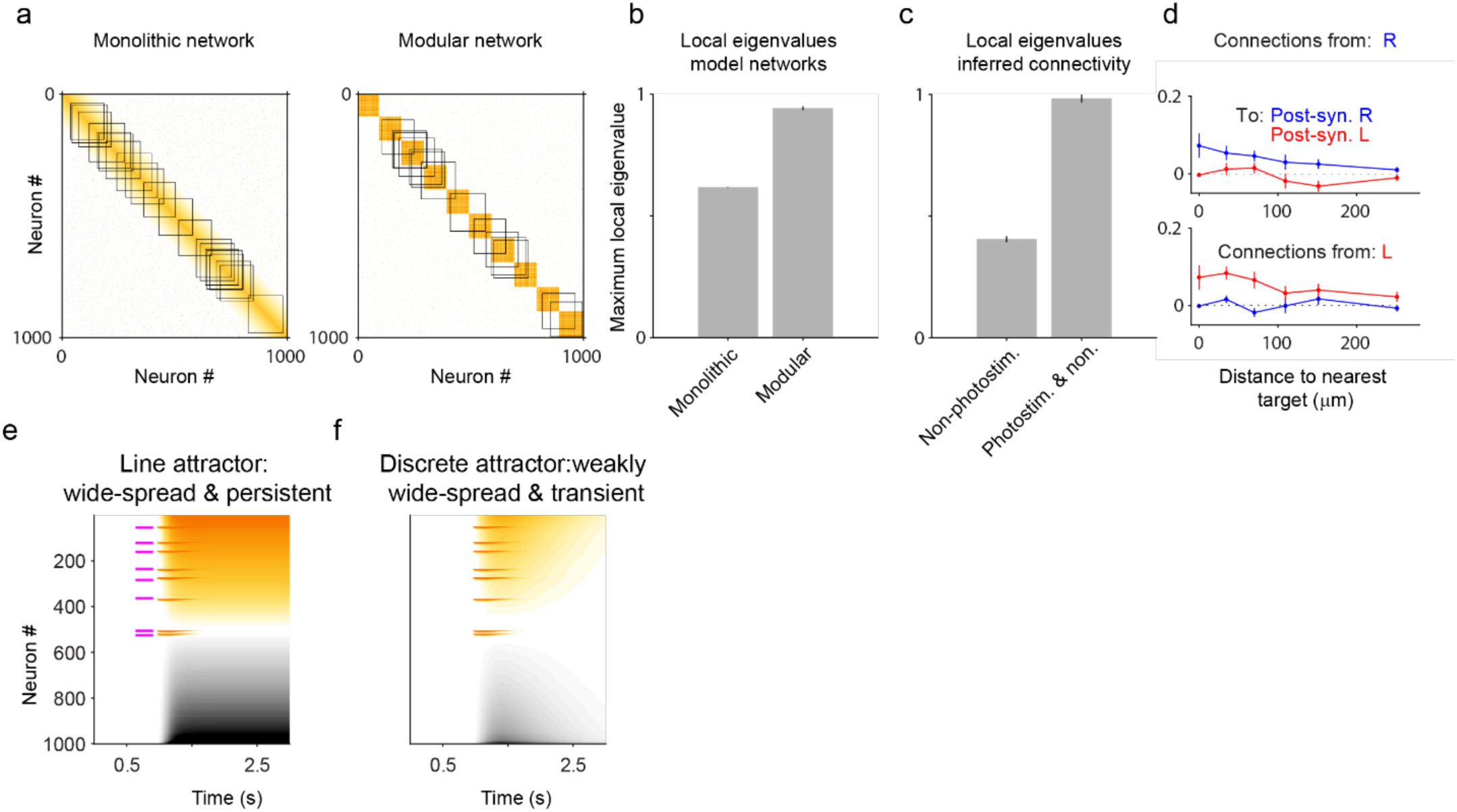
The key difference between the two model networks with locally biased connectivity in Figure 4b-c is the presence of a single persistent mode of activity in the monolithic network and multiple distinct persistent modes in the modular network. More precisely, the monolithic network has a single mode with eigenvalue equal to 1 and the modular network has 10 such eigenvalues. We use this fact to develop an analysis aimed at determining which of these two models more closely resembles the connectivity inferred from the data in Figure 4d-e. **a**. If we were to draw a box around one of the modules in Figure 4c, we would find that the maximum eigenvalue of this subnetwork is 1, whereas an equivalent box in the monolithic model in Figure 4b would have a maximum eigenvalue of 0.48. To extend this “local eigenvalue” analysis to the inferred networks in Figure 4d-e we wouldn’t know which neurons to draw this box around, so to make the analysis more general, we instead draw many boxes at random locations along the diagonal and calculate the maximum eigenvalue of subnetwork within the box. **b**. On average, the modular network has larger local eigenvalues (mean, 0.94, range, 0.84 — 1.0, 75% CI) than the monolithic network (mean, 0.62, range, 0.61 — 63, 75% CI; error bars, s.e.m.). **c.** For the inferred connectivity matrix (Fig. 4d-e), we identify all neurons within 70 µm of a photostimulation target whose activity was significantly perturbed by photostimulation (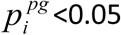, methods), and calculate the maximum eigenvalue of this subnetwork. We find that the largest local eigenvalue in networks trained to match photostimulation and non-photostimulation trials was similar to the modular network in Fig. 4c (mean, 0.98, range 0.75 — 1.27, 75% CI). Networks trained to match the activity during non-photostimulation trials only (**d**) had a maximum eigenvalue that were similar to the monolithic network (mean, 0.41, range, 0.19 — 0.62, 75% CI). **e-f.** Response of a line attractor (**e**) and discrete attractor (**f**) network (Methods) to targeted photostimulation of 8 neurons. Due to widespread connectivity, perturbations of a small number of neurons spread throughout both networks.

## Methods

### Mice

Data are from eleven mice (age at the beginning of experiments, 70 – 150 days). Eight mice were used for behavioral experiments. Two mice were used for characterization of the spatial resolution of photostimulation (Fig1. c, Extended Data Fig. 6) and one mouse for electrophysiology experiments (Extended Data Fig. 5). All mice were from the CamK2a-tTA (JAX:007004) x Ai94(TITL-GCaMP6s)^1^ (JAX: 024104) transgenic line. Half of the mice were crossed with Emx1-Cre (JAX: 005628) and the other half were crossed with slc17a7 IRES Cre (JAX: 023527), both providing broad expression of GCaMP6s in excitatory cortical neurons.

### Surgical procedures

All procedures were in accordance with protocols approved by the Janelia Institutional Animal Care and Use Committee. During surgical procedures mice were anesthetized using 1-2 % Isoflurane. After surgery mice were given Buprenorphine HCl (0.1 ml, 0.03 mg/ml). Ketoprofen (0.1 ml, 0.1 mg/ml) was provided on the day of surgery and for two days following surgery. Three millimeter circular craniotomies were centered over ALM (2.5 mm anterior and 1.5 mm lateral from Bregma). Virus (10^12^ titer; AAV2/2 camKII-KV2.1-ChrimsonR-FusionRed; Addgene, plasmid #102771) was injected 400 μm below the dura (4-10 sites, 20-30 nl each; or 2 sites, 100 nl each), centered within the craniotomy and spaced by 500 μm. The craniotomy was covered by a cranial window composed of 3 layers of circular glass (total thickness, 450 μm). The diameter of the bottom two layers was 2.5 mm diameter. The top layer was 3 mm or 3.5 mm and rested on the skull. The window was cemented in place using cyanoacrylate glue and dental acrylic (Lang Dental). A custom headbar was attached just anterior to the window using cyanoacrylate glue and dental cement. After 3-7 days of recovery mice were placed on water restriction (1 ml/day). Behavioral training started 3-5 days later. Coexpression of GCaMP6s and ChrimsonR was confirmed in histological sections imaged using an inverted confocal microscope (Extended Data Fig. 1; Zeiss, LSM 880 Airyscan).

### Behavior

Behavioral training was performed as described previously^2,3^. Briefly, mice were presented with one of two auditory cues. Half of the mice were trained to discriminate between 3kHz and 12kHz pure tones and the other half were trained to discriminate between white noise or an equally weighted combination of pure tones with frequencies of 0.5, 1, 2, 4, 8 and 16 kHz. No qualitative differences in ALM activity resulted from using different sets of auditory stimuli. During the sample epoch (1.25 seconds) white noise was played continuously whereas tones were played for five repetitions of 150 ms pulses with 100 ms inter-pulse intervals. During the sample and delay epochs, the lickports were out of reach of the mouse^4^. The lickports began moving towards the mouse 2.6 seconds after the start of the delay, arriving 3 seconds after the start of the delay. The arrival of the lickport served as a ‘go cue’ for the mice to begin licking. Mice were allowed to lick for reward 3 seconds after the start of the delay epoch.

Imaging and photostimulation experiments were started once the mice achieved performance of > 65%, typically 4-6 weeks after the start of training. Statistical power analysis revealed that 65 % performance maximizes our ability to detect changes in behavior caused by photostimulation. An exploratory round of experiments (3 mice, 33 sessions), in which 4 or 5 groups were photostimulated per experiment, showed small but robust changes in mouse behavior after photostimulation. Based on these preliminary experiments, in the second round of experiments we enhanced our ability to detect changes in behavior by increasing the number of trials per photostimulation group (5 mice, 51 sessions) at the expense of the number of photostimulation groups (two per session). Given the number of photostimulation trials per session (mean, 25; range, 13 — 38, 75% CI; per trial type), statistical power analysis indicated that behavioral changes greater than ± 18% correspond to p < 0.05 in single sessions.

### Microscope

Two-photon imaging and two-photon photostimulation were performed using a custom microscope^4^ with an additional of a photostimulation path, consisting of a 1040 nm pulsed laser (Fidelity 10, Coherent), a Pockels cell (Conoptics) for power modulation, and a pair of galvanometer mirrors (Cambridge, 6215H) for beam positioning (Extended Data Fig. 2). Imaging was with 920 nm light (Chameleon Ultra II, Coherent) and a resonant scanner (Thorlabs). Imaging and photostimulation were controlled by Scanimage 2016a (Vidrio).

The field of view was adjusted according to the spatial range of opsin expression (one of: 585μm x 599μm, 512×512 pixels, 30 Hz; 814μm x 732μm, 640×640 pixels, 24 Hz; 968μm x 822μm, 700×700 pixels, 22 Hz). Imaging was restricted to layer 2/3 of anterior lateral motor cortex (ALM), 125-250 microns below the surface of the brain.

### Photostimulation and behavior experiments

Neurons were chosen for membership in a photostimulation group based on their selectivity in either the early delay, late delay or early response epochs. Selectivity of individual neurons was either determined based on activity in the first 30-70 trials in a session, or activity measured on the previous day. After defining 2-5 photostimulation groups (8 neurons each) for a session, the photostimulation targets were loaded into ScanImage using custom Matlab software. Each neuron was photostimulated for 3 ms each, with 1 ms between photostimulation of different neurons (Extended Data Fig. 4). After all 8 neurons had been photostimulated (8x(3ms+1ms)=32ms) the first neuron was photostimulated again until each neuron was photostimulated 10 times, for a total of 10×32ms = 320ms. The power of the photostimulation beam at the sample was 100-150mW. Because this power was sufficient to excite GCaMP fluorescence, all imaging frames acquired during the photostimulation were discarded. Photostimuli were typically provided one second after start of the delay epoch (138/215 photostimulation groups), but in some experiments we photostimulated one second earlier (61/215 groups) or later (16/215 groups). Photostimulation and control trials were randomly interleaved, with photostimuli delivered on either 33% or 40% of trials.

### Electrophysiology

Extracellular voltage was recorded in lightly anesthetized mice (0.75% isoflurane) with cell-attached recordings. Signals were acquired using an Axopatch 700B amplifier (Molecular Devices) at 20 kHz (http://wavesurfer.janelia.org). Electrodes with 10 MΩ impedance were filled with ACSF (in mM): 125 NaCl, 5 KCl, 10 dextrose, 10 HEPES, 2 CaCl_2_, 2 MgSO_4_, pH 7.4 and advanced toward a neuron of interest in layer 2/3 of primary motor cortex with 1 psi of positive pressure. Neurons were photostimulated with a train of 10 3-ms spirals with a 28-ms inter-spiral interval at 50, 100 and 150 mW.

### Analysis of calcium-related fluorescence dynamics

Regions of interest (ROIs) corresponding to cell bodies were generated using a semi-automated algorithm^5^. Cell bodies were identified in images averaged across a session, and images in which all frames corresponding to correct lick-left trials were averaged and subtracted from an average of all frames from correct lick-right trials. These selectivity maps (Extended Data Fig. 3a) were used for choosing neurons for photostimulation and also for ensuring that all selective neurons were included in analysis. *f* (*t*) = Δ*F*(*t*)/ *F*_0_ was calculated for each cell by defining baseline fluorescence (F_0_) as the average fluorescence during the pre-sample period averaged across all trials. Fluorescence traces were then separated by trial-type. The fluorescence trace at time *t* of neuron *i* on trial number *j*, trial type *k* (left, k = L; right, k = R; both left and right, k = L&R) and photostimulation group number *s* (non-photostimulation, s = non.; photostimulation of group *pg, s* = *pg*) is 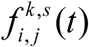. In this notation the fluorescence of neuron *i* on the *jth* lick right trial with photostimulation of group *pg* is 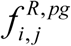.

### Selectivity

Selectivity for trial type was used to choose neurons for photostimulation and to group neurons for analysis. Selectivity *S*_*i*_ is the trial-averaged difference between fluorescence for left and right correct non photostimulation trials, around the go cue (*t*_*cue*_):

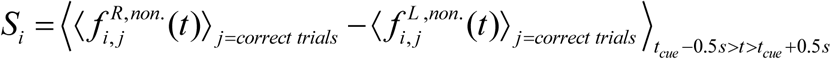

### Change in activity caused by photostimulation

The trial-averaged change in activity produced by the photostimulation group *pg* on the *ith* neuron is (Fig. 1f,h):

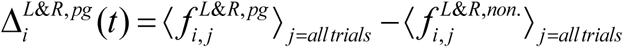

To characterize the strength of photostimulation in each neuron and trial (Fig. 1g) we compared the time-averaged fluorescence with and without photostimulation:

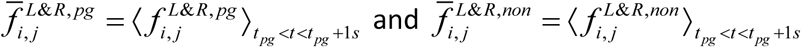

where *t*_*pg*_ is the end of the photostimulus for group 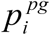 is the p-value (two-tailed t-test) comparing the distributions of 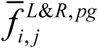 and 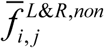 for each neuron. We refer to all neurons that were within 20 µm of a photostimulation target with 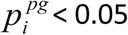 as directly photostimulated. Neurons with 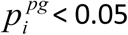 were plotted in Figure 1g (‘perturbed’).

The average selectivity of all directly photostimulated neurons is 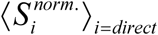 (Fig. 1f, gray shading; Fig. 1j; Fig. 2b, x-axis; Fig. 2c, bottom, y-axis). Here 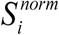 is the average selectivity, *Si*, divided by the standard deviation in fluorescence of neuron *i* (z-scored), normalized to have R^2^ norm equal to one.

### Persistence of photostimulation

To quantify the persistence of the change in activity produced by photostimulation, we averaged the trial-averaged change in activity across all directly photostimulated neurons,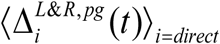 (Figure 1h). We define the decay time constant *τ* _*pg*_ as the time at which 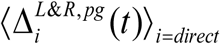 decays to 1/e of its peak value. Because many photostimulation groups produce changes that remain larger than 1/e of the peak after 3 seconds, we fit 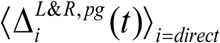 using

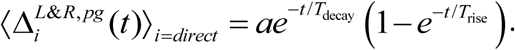

*τ* _*pg*_ is the time when the fit decayed to 1/e (Figure 1h-j). Only photostimulation groups with photostimulation starting at one or zero seconds after the start of the delay epoch (199/215 groups) were included in analysis of persistence. In Figure 1 i-j data were binned in quintiles based on their value along the x-axis.

### Relationship between directly photostimulated neurons and coupled neurons

The coupling strength from directly photostimulated neurons in group *pg* with neuron *i* was calculated as:

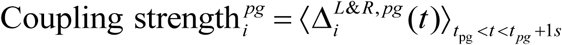

for all neurons more then 30 µm from a photostimulation target (Fig. 2b-d, y-axes). Coupling in R (Δ*R*^*pg*^) and L (Δ*L*^*pg*^) populations was calculated as the average of the coupling strength for all R and L (Fig. 2b, y-axis) neurons where the contribution of each neuron was weighted by the normalized selectivity 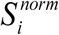:

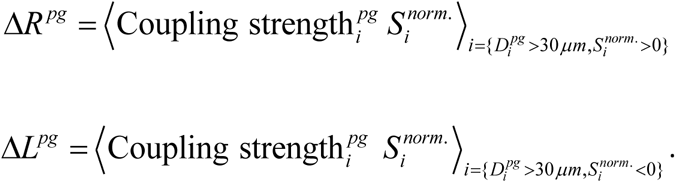

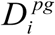 is the distance of neuron *i* to the nearest photostimulation target in group *pg*.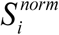 and Coupling 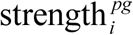 were computed by using non-overlapping groups of randomly sampled non-photostimulation trials. Data were binned in quintiles according to the selectivity of the directly photostimulated neurons (Fig. 2b, x-axis). We further analyzed coupled responses to photostimulation of the groups with the strongest right selectivity (Top quintile; Fig. 2b, dashed box). To assess the distance dependence of specific coupling, we calculated Δ*R*^*pg*^ and Δ*L*^*pg*^ in bins of 20 µm width from 30 —250 µm (Fig. 2c, top). In these bins we also calculated the average normalized selectivity 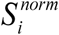 (Fig. 2c, bottom).

We calculated the noise correlation (Fig. 2d) between two neurons *i* and *j* as:

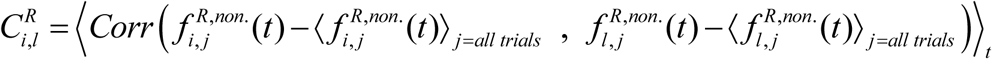

The average correlation of coupled neuron *i* with photostimulation group *pg* was 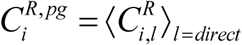. A similar analysis was performed for left trials. Noise correlations for right and left trials were then averaged (Fig. 2d, x-axis):

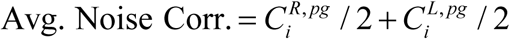

### Changes in behavioral performance

For each photostimulation group, we calculated the difference in correct response rate between photostimulation trials and non-photostimulation trials separately for lick left and lick right trials. P values were calculated by downsampling the non-photostimulation trials to match the number of photostimulation trials. This process was repeated 10,000 times with replacement (Figure 3a, Extended Data Fig. 8). To estimate the false-positive rate, we calculated 10,000 null-distributions of p-values from the downsampled non-photostimulation trials (Figure 3b, gray line) and compared this to the distribution of p-values from photostimulation trials (Figure 3b, black line) using the Kolmogorov-Smirnov test.

To relate changes in behavior to photostimulated changes in activity, we calculated ‘activated population selectivity’ (Fig. 3c-f) as the overlap between the photostimulated change in activity 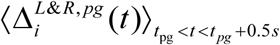 and trial-type selectivity, *S*_*i*_. At the population level, the change in activity induced by photostimulation was relatively sparse (Fig. 1d, Fig. 4a) and trial-to-trial variability was substantial. We isolated the effects mediated by photostimulation from normal trial-to-trial variability. We first divided the photostimulation trials into two halves, a testing and training set. We next determined a photostimulation subspace **V**_*pg*_ via an SVD decomposition on the first half of the photostimulated data for time points immediately following photostimulation:

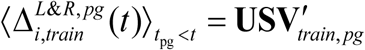

We then projected the change in activity from a separate subset of trials 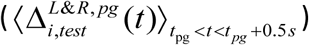 onto this photostimulation subspace to obtain the photostimulation vector (Fig. 3c-e).

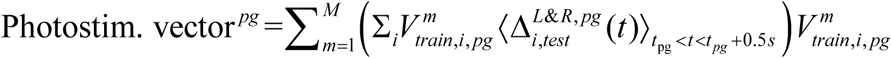

where the *m* denotes the *mth* column of **V**_*train, pg*_. Because we found that 93% of the RMS signal of 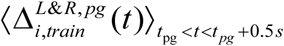 was contained in the first mode of **V**_*pg*_ we limited analysis to this mode (i.e. M=1). A qualitatively similar relationship between activated population selectivity and behavioral bias was observed for M = 2. Activated population selectivity (Fig. 3f; x-axis) was calculated as:

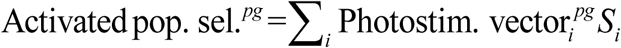

Random sampling of training and testing subsets was repeated 10 times and the average activated population selectivity was computed for each photostimulation group. The same analysis was repeated by splitting non-photostimulation trials into two non-overlapping sets and treating one set as if it were from a photostimulation experiment. This sham photostimulation subset was selected to have the same number of trials as the actual photostimulation experiment (Fig. 3f, gray), and was further split in half into randomly sampled training and testing subsets for 10 repetitions as described above for the actual photostimulation experiments.

#### Network models

The spike rate of model neuron *i* was determined by the equation:

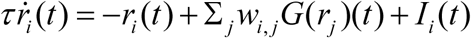

Where *w*_*i,j*_ is the connection from neuron *j* onto neuron *i, τ*is the synaptic time constant (taken to be 100 ms), *G* is the synaptic non-linearity and *I*_*i*_ (*t*) is the external input. For the non-modular network with local connectivity (Fig. 4b) the connectivity was *w*_*i,j*_ = *e*^−|*i*– *j*|^/^70^. For the monolithic attractor models (where all neurons contribute to the attractor) (Extended Data Fig. 10a, b) *w* was a rank-one matrix constructed by taking the outer product of a Gaussian random 1000×1 norm-1 vector with itself. For the line-attractor model (Extended Data Fig. 10a), the synaptic function was *G*(*x*) = *x*; the discrete attractor model (Extended Data Fig. 10b) had *G*(*x*) = 0.1tanh(0.1*x*). The modular attractor network (Fig. 4c) consisted of 10 rank-one line-attractor networks, each composed of 100 neurons and *G*(*x*) = *x*. Intermodular connections were sparse, with 96% of connections set to zero and the remaining connections drawn randomly from a Gaussian distribution with mean zero. Short-term memory was simulated by giving a brief input to each network along the network’s primary eigenvector. Photostimulation was simulated by injecting a brief current to 8 randomly chosen neurons.

#### Generating model connectivity by fitting to data

Connectivity matrices were trained to fit the fluorescence activity of all individual neurons^6^. For each experimental session with *N* recorded neurons we fit the *N*x*N* matrix *w*_*i,j*_ to reproduce the fluorescence of each neuron. To account for the time-course of calcium-sensitive fluorescence we approximated the network equation to be:

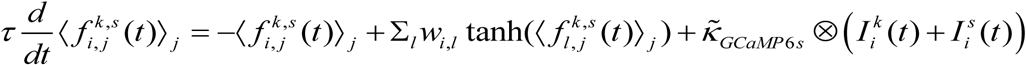

We used the approximate kernel 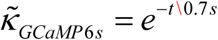, which reflects the slow decay component of the GCaMP6s response to a burst of spikes (Extended Data Fig. 7). External sensory 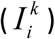 and photostimulation 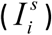 currents were step functions that were only active during the sample and photostimulation epochs, respectively. Fitting was done using a recursive least-square algorithm^6,7^ in which all weights onto a given neuron are tuned at each time step to minimize the difference between its modeled and experimentally observed fluorescence activity. Activity from different trial types (k = left & k = right) and photostimulation conditions (s = non., s = pg 1 … pg N) were fit sequentially. Each fit was iterated 30 times, which produced high-quality fits (Median Pearson correlation = 0.69, range 0.25—0.92, 75% CI). For each session, two separate fits were obtained, one to capture only non-photostimulation trials and the other to capture both photostimulation and non-photostimulation trials.

